# “Myosin XI drives polarized growth by vesicle clustering and local enrichment of F-actin in *Physcomitrium (Physcomitrella) patens*”

**DOI:** 10.1101/2020.08.25.266296

**Authors:** Giulia Galotto, Pattipong Wisanpitayakorn, Jeffrey P. Bibeau, Yen-Chun Liu, Parker J. Simpson, Erkan Tüzel, Luis Vidali

## Abstract

In tip-growing plant cells, growth results from myosin XI and F-actin mediated deposition of cell wall polysaccharides contained in secretory vesicles. Previous evidence showed that myosin XI anticipates F-actin accumulation at the cell’s tip, suggesting a mechanism where vesicle clustering via myosin XI promotes F-actin polymerization. To evaluate this model, we used a conditional loss-of-function strategy by generating *Physcomitrium (Physcomitrella) patens* plants harboring a myosin XI temperature-sensitive allele. We found that loss of myosin XI function alters tip cell morphology, vacuolar homeostasis, and cell viability, but not following F-actin depolymerization. Importantly, our conditional loss-of-function analysis shows that myosin XI clusters and directs vesicles at the tip of the cell, which induces F-actin polymerization, increasing F-actin’s local concentration. Our findings support the role of myosin XI in vesicle clustering and F-actin organization, necessary for tip growth, and deepen our understanding of additional myosin XI functions.

## INTRODUCTION

Myosin XI is an essential plant molecular motor involved in vesicular transport, organelle motility, and plant growth (Tominaga et al., 2003; Peremyslov et al., 2008; Madison et al., 2015; Abu-Abied et al., 2018; Duan and Tominaga, 2018; Ryan and Nebenfuhr, 2018)}. In tip growing cells, it is assumed myosin XI mediates growth via the active transport of secretory vesicles dependent on the F-actin cytoskeleton (Ojangu et al., 2007; Peremyslov et al., 2008; Vidali et al., 2010; Park and Nebenfuhr, 2013; Bibeau et al., 2020). This is the case for pollen tubes and root hairs in flowering and vascular plants and protonemal cells in bryophytes, which grow in a highly anisotropic fashion. In these cells, secretory vesicles containing cell wall polymers and wall loosening enzymes are delivered to the apex, where their secretion is locally regulated by Rac-Rop GTPases (Hepler et al., 2001; Kost, 2008; Rounds and Bezanilla, 2013; Le Bail et al., 2019). In the area of vesicle fusion, newly generated cell wall has enhanced extensibility; for the cell to expand without bursting and to reach fast growth rates (5.5 nm/sec in *Physcomitrium (Physcomitrella) patens* protonemata and 200-300 nm/sec in *Lilium longiflorum* pollen tubes), the growth mechanism needs to be tightly regulated. The motor myosin XI and the F-actin cytoskeleton are the main components of the growth machinery. In the past decades, an extensive amount of research has been performed to understand how a variety of cellular processes interact to generate and maintain such an anisotropic growth fashion (Cole and Fowler, 2006; Kroeger and Geitmann, 2012; Rounds and Bezanilla, 2013; Orr et al., 2020). Despite extensive research, however, this phenomenon is not completely understood. Multiple lines of evidence suggest myosin XI is key to the development of proper cell polarization (Tominaga et al., 2000; Ojangu et al., 2007; Peremyslov et al., 2008; Peremyslov et al., 2010; Vidali et al., 2010; Cai et al., 2014). In land plants, such as *P. patens*, a knockdown approach showed myosin XI is essential for maintaining polarized growth (Vidali et al., 2010). In the vascular plant *Arabidopsis thaliana*, myosins XI (a class with 13 genes) is involved in the development of root hairs and trichome morphogenesis, as well as organelle trafficking (Ojangu et al., 2007; Peremyslov et al., 2008). Some reports show myosin XI is also important for F-actin organization; such is the case in epidermal *A. thaliana* cells (Peremyslov et al., 2010; Cai et al., 2014) and root hairs (Tominaga et al., 2000). The F-actin cytoskeleton, together with myosin XI, plays a crucial role in tip growth. In *L. longiflorum* pollen tubes, depolymerization of F-actin through latrunculin B, Cytochalasin D, and DNAseI, stops growth (Vidali et al., 2001; Finka et al., 2007). In *L. longiflorum* pollen tubes, F-actin polymerization is a limiting factor for pollen tube elongation, in a process independent from cytoplasmic streaming (Vidali et al., 2001). Similarly to pollen tubes, the pharmacological inhibition of F-actin in *P. patens* protonemata alters F-actin structures and stops the growth in a dose-dependent manner (Vidali et al., 2009a). Furthermore, in *P. patens*, an active transport that relies on F-actin is essential for tip growth: a mechanism based on vesicle diffusion alone cannot support the observed growth rates (Bibeau et al., 2018).

Fluorescence cross-correlation analyses of F-actin, myosin XI, and VAMP-labeled vesicles in *P. patens* characterized their temporal relation in protonemal cell tips (Furt et al., 2013). Surprisingly, while myosin XI and the VAMP-labeled vesicles are in phase, the myosin XI and the actin signal are not, with the myosin XI-vesicle signal leading the F-actin signal by ~18 sec (Furt et al., 2013). This is surprising because a transport mechanism based on myosin XI motors walking on preassembled F-actin would result in the F-actin signal leading or being temporally correlated with the myosin XI signal. Instead, the myosin XI-vesicle cluster leading the F-actin suggests a different cellular mechanism than the one provided by the simple F-actin track hypothesis.

In this work, we hypothesize myosin XI clusters vesicles enabling local F-actin polymerization, and that this mechanism is essential for the maintenance of apical growth. We tested our hypothesis by generating a temperature-sensitive (TS) allele of myosin XI and by combining it with the vesicle marker 3mCherry-VAMP and the F-actin marker Lifeact-mEGFP. The generation of loss of function via TS alleles has been paramount to uncover the role of essential genes in plants, but have been historically limited to forward genetic screens (Lane et al., 2001; Whittington et al., 2001). Here, we generated a conditional myosin XI allele through rational mutagenesis. We used the plant *P. patens*, which is a great model to study polarized growth due to its simple cytology and tractable genetics (Reski, 2018; Rensing et al., 2020). Our results prove that myosin XI clusters secretory vesicles at the cell’s apex and that myosin XI is essential for persistent apical growth. In addition, we show that myosin XI is necessary for the preservation of the elongated cell morphology and proper apical wall curvature, characteristic of tip growing cells. Finally, we also observed myosin XI is involved in vacuole homeostasis and cell viability.

## RESULTS

### Development of a myosin XI temperature-sensitive allele

Myosin XI function is essential for tip growth (Vidali et al., 2010); hence, to be able to further characterize its function in polarized growth, we generated a moss line harboring a TS allele of myosin XI. To do so, we used a previously described strategy to generate temperature-sensitive alleles in plants (Vidali et al., 2009b). This strategy is based on mutagenizing residues in the core of the protein of interest, which will render it less stable at higher temperatures, likely resulting in protein unfolding or denaturation (Vidali et al., 2009b). We selected potential buried residues to target by assigning a hydrophobicity score to each amino acid in the myosin XI protein sequence (Kyte and Doolittle, 1982); then, we identified the ones that confer temperature sensitivity to the plants (20°C vs. 32°C) via an RNAi-based complementation assay (Vidali et al., 2007) (see Methods). In the *P. patens* genome, there are two myosin XI genes, myosin XIa and myosin XIb, which are 94% identical at the protein level. Here we decided to modify myosin XIa because it is expressed at higher levels (Vidali et al., 2010). We found that we needed to insert two point mutations in the myosin XIa gene: V584A, L616A to render the protein temperature-sensitive (Figure 1a,b, and Methods). Due to the functional redundancy of the two myosin XI genes (Vidali et al., 2010), we coupled the site-directed mutagenesis with the deletion of myosin XIb (line referred to as myoXIaTS). As a control for the loss of myosin XIb, we generated a *P. patens* line maintaining the WT version of myosin XIa, but harboring the deletion of myosin XIb (line referred to as myoXIaWT). Both lines were generated in a parental line expressing Lifeact-mEGFP (Vidali et al., 2009a), which was also used as a control.

**Figure 1.**
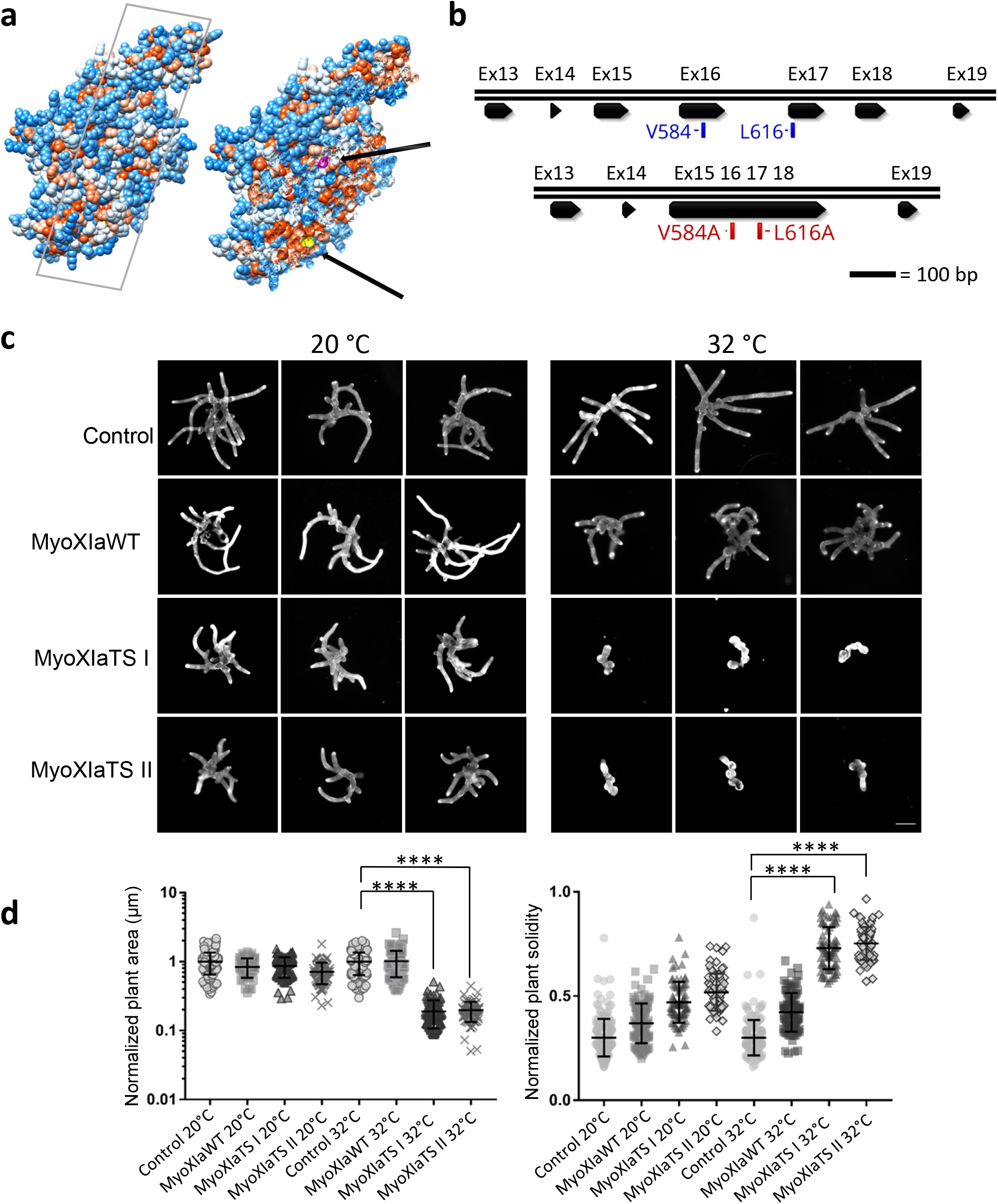
Two point mutations (V584A, L616A) render myosin XI protein temperature sensitive. **a.** Myosin XIa motor domain modeled on the crystal structure of Myosin Vc from *Homo Sapiens* with Swissmodel. The amino acid hydrophobicity was color-coded on the Kyte-Doolittle scale; low hydrophobicity is depicted in blue, high in orange-red. The surface of the domain was sliced to show the two point-mutations buried in the hydrophobic core, Valine in yellow and Leucine in Magenta (arrows). **b.** Myosin XIa TS, and Myosin XIb KO. **c.** Representative images of 1-week old control and mutant plants grown at 20°C and 32°C. The plants labeled as “control” are the parental line control. In the panel, the cellulose is stained with calcofluor-white. Myosin XIa TS I and II represent two independent lines harboring the temperature-sensitive allele. Scale bar = 50 μm. **d.** Quantification of normalized plant area and solidity of control and mutant plants at grown at 20°C and 32°C. Number of plants analyzed: control 20°C, 174. Control 32°C, 163. Myosin XIb KO 20°C, 96. Myosin XIb KO 32°C, 92. Myosin XIb KO Myosin XIa TS I 20°C, 76. Myosin XIb KO Myosin XIa TS I 32°C, 78. Myosin XIb KO Myosin XIa TS II 20°C, 80. Myosin XIb KO Myosin XIa TS II 32°C, 73. Error bars represent the standard deviation of the mean. Comparison among groups performed via 2-way ANOVA (adjusted *P*< 0.0001).

To confirm the effect of the myosin XIaTS protein on plant growth and morphology, we performed a morphometric assay (Galotto et al., 2019). After regeneration of protoplasts for four days, plants were exposed at 20°C and at 32°C for three days, then imaged. When plants are lacking myosin XIb but express the WT copy of myosin XIa, they do not exhibit morphological defects: they appear polarized, and the plant area is comparable to the parental line (Figure 1c). On the other hand, plants expressing the TS allele of myosin XIa and lacking the functional copy of myosin XIb, exhibit strong temperature sensitivity at 32°C (Figure 1c). Mutant plants at 32°C do not have tip growing cells and are highly stunted (Figure 1c). Plant area and solidity were computed to quantify the observed differences (Vidali et al., 2010; Galotto et al., 2019) (Figure 1d). The plant area is comparable in controls at the permissive temperature, and it is significantly reduced in TS plants at the restrictive temperature. Solidity values are low in control plants at 32°C and are closer to 1 in TS plants at 32°C, characteristic of round and unbranched plants, and consistent with previous RNAi results (Vidali et al., 2010). These data show that plants expressing the mutant (V584A, L616A) allele of myosin XIa and lacking a functional copy of myosin XIb are temperature sensitive at 32°C.

### Myosin XI loss-of-function causes cell death while simultaneous F-actin depletion restores viability

During preliminary experiments aimed to characterize the myoXIaTS line, we observed that, when exposed at 32°C for 24 hrs, many cells in the plant appeared dead. To further investigate if prolonged lack of functional myosin XI leads to cell death, we incubated myoXIaWT and myoXIaTS cells at 20°C and 32°C for 24 hrs. Prior to imaging, plants were stained with calcofluor-white and propidium iodide, to mark the plant’s outline and the nucleus of dead cells, respectively. After 24 hrs of incubation at 32°C, a high number of myoXIaTS apical cells die, compared to the myoXIaWT plants at 32°C (Figure 2). In contrast, in the myoXIaWT cells exposed at 32°C only a few cells die, probably due to temperature stress.

**Figure 2.**
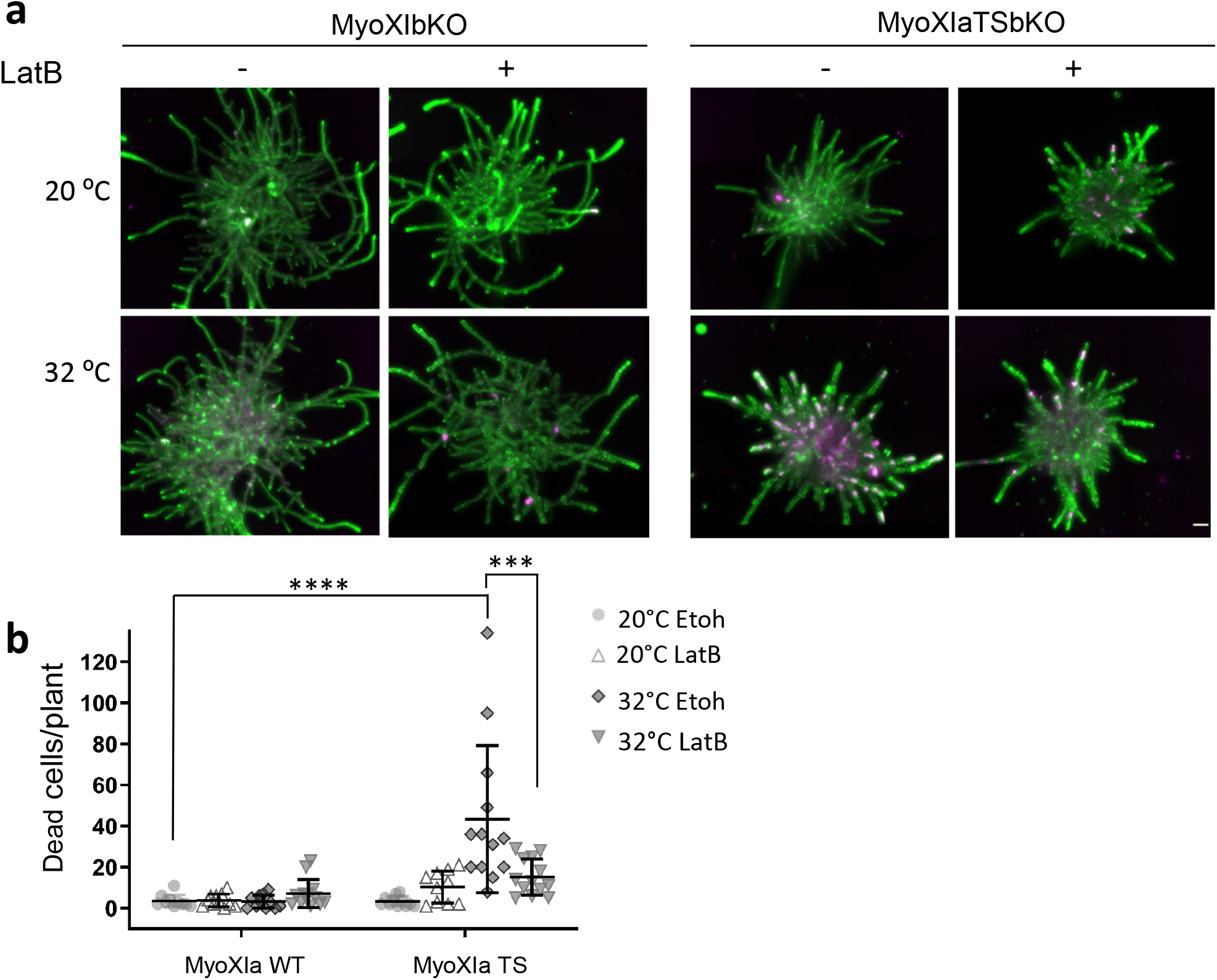
Myosin XI TS cells dye after 24 hrs exposure at 32°C, plants, depolymerization of F-actin via Latrunculin B reverts the phenotype. **a.** Representative control and myosin XIa TS cells exposed at 32°C for 24 hrs, with ethanol (vehicle control) or Latrunculin B (20 μM). In the panel, green represents calcofluor staining and magenta the dead cells stained with propidium iodide. **b.** Quantification of the number of dead cells per plant. Number of plants analyzed: control 20°C Etoh, 11. Control 20°C LatB, 11. Control 32°C Etoh, 11. Control 32°C LatB, 11. Myosin XIa TS 20°C Etoh, 11. Myosin XIa TS 20°C LatB, 11. Myosin XIa TS 32°C Etoh, 11. Myosin XIa TS 32°C LatB, 11. Differences between groups are computed via a 3 way ANOVA (adjusted *P*< 0.0001). Error bars represent the standard deviation from the mean. Scale bar = 100 μm.

Since myosin XI is a F-actin dependent motor, we decided to investigate whether, in a similar way to myosin XI loss-of-function, F-actin depolymerization affects cell survival at 32°C. To test this, we treated both cell lines at 20°C and 32°C with 20 ◻M latrunculin B, which fully depolymerizes F-actin (Vidali et al., 2009a; Bibeau et al., 2020). Surprisingly, when myoXIaTS plants were exposed to 32°C and simultaneously treated with latrunculin B, cell viability was rescued (Figure 2). Consistent with this, in the latrunculin B treated myoXIaWT plants, only a very small number of cells died, either at 20°C or 32°C (Figure 2). These results show that the inactivation of myosin XI in cells in which the F-actin cytoskeleton is still present strongly affects cell survival and are consistent with the hypothesis that myosin XI is essential for F-actin organization in protonemata. To get insights into the molecular mechanism triggering the observed cell death, we performed preliminary RNASeq on control and myoXIaTS plants exposed at 32°C for 2 hours. Differential expression analysis of these preliminary results suggests that cell death is not due to the activation of a programmed cell death pathway. Further experiments will be needed to verify if a programmed cell death pathway is activated at time points later than 2 hours.

### In cells with decreased myosin XI activity, but not after F-actin depletion, the vacuole dilates and invades the cell tip

While observing myoXIaTS cells growing at 32°C, we noticed the expansion of the vacuole at the tip of the cell. Vacuoles are essential plant organelles involved in the preservation of cell homeostasis and the support of cytoplasmic turgor pressure, essential for cell shape establishment and maintenance (Marty, 1999). The vacuole in *P. patens* protonemal tip cells reflects the characteristic “zonation,” where an expanded section of the vacuole occupies the back of the cell, while a tubular anastomosing section of the vacuole populates the apical region of the cell (Oda et al., 2009; Furt et al., 2012). To investigate the dynamics of the observed vacuolar alterations in the myosin XI depleted plants, we incubated myoXIaWT cells and myoXIaTS cells at 20°C and 32°C for 2 hours, and we stained them with the tonoplast marker MDY-64 (Scheuring et al., 2015) (Figure 3). Control cells exhibit, as expected (both at 20°C and 32°C), a high number of anastomosing sections of the vacuole, especially in the region between the nucleus and the sub-apex (Figure 3). However, in the myoXIaTS cells incubated at 32°C for 2 hours, we observed the disappearance of the anastomosing sections, and instead, the vacuole enlarges and invades the tip (Figure 3). As performed in the previous experiment (Figure 2), we further investigated whether the depolymerization of F-actin by latrunculin B (20 μM) would mimic the myoXIaTS vacuolar phenotype. To our surprise, F-actin depolymerization did not phenocopy the TS phenotype, but instead, the MyoXIaTS cells at 32°C and treated with latrunculin B present a highly anastomosed vacuole, similar to controls. These results indicate that myosin XI function is important for vacuolar homeostasis and structure. Interestingly, these results are similar to the results from the cell death assay, in that when F-actin is depleted, the lack of myosin XI has reduced effect on the cells. Together, these results are consistent with the hypothesis that myosin XI participates in F-actin organization during growth.

**Figure 3.**
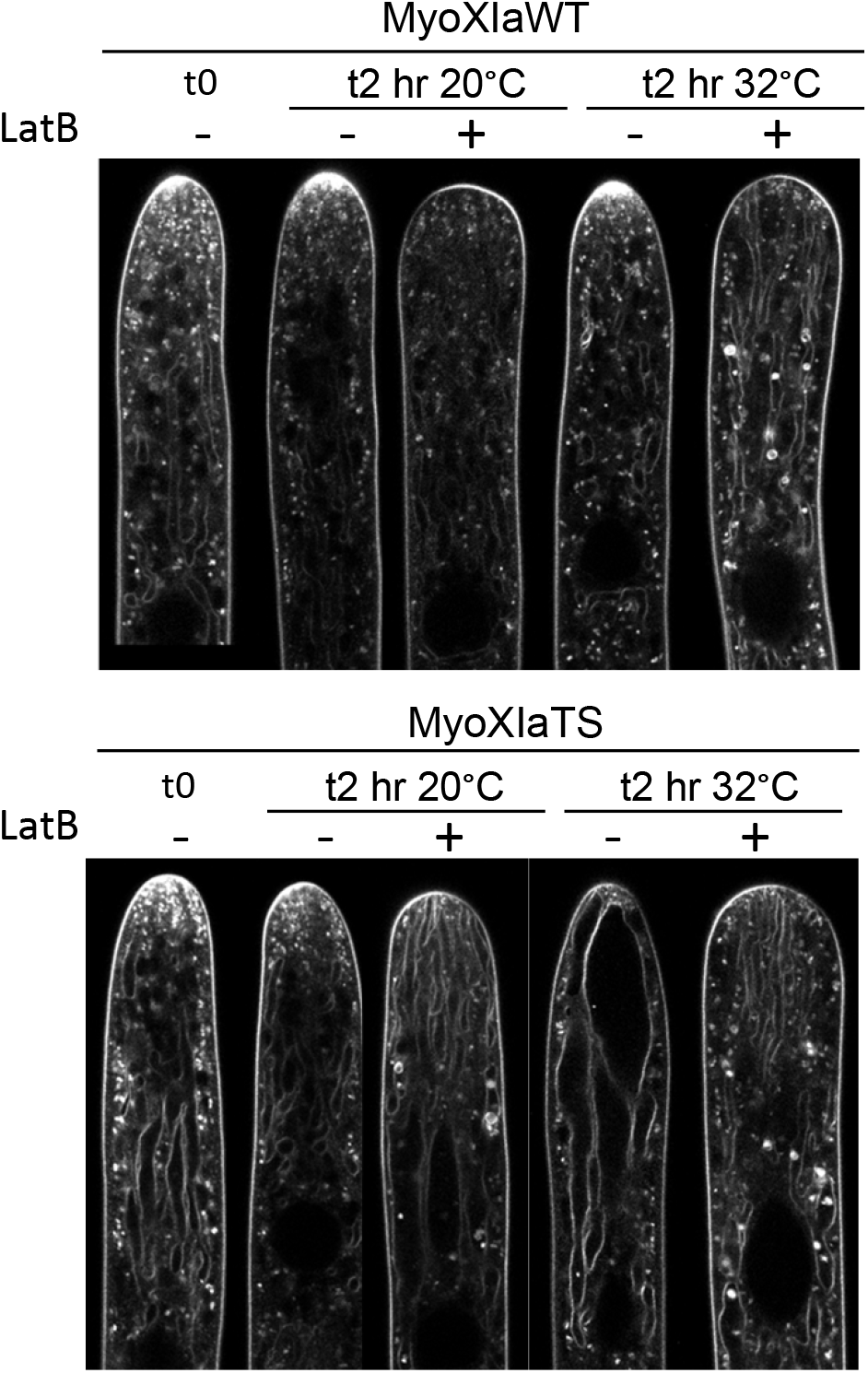
Myosin XIa TS cells present an enlarged aberrant vacuole, depolymerization of F-actin via latrunculin B reverts the phenotype. Representative control and myosin XIa TS cells exposed at 32°C for 2 hrs, with ethanol (vehicle control) or latrunculin B (20 μM). The tonoplast is stained with MDY-64 (500 nM) Note that the dye also stains the plasma membrane and membranous organelle within the cell. For display purposes, the images were γ-adjusted (0.8%). Scale bar = 10 μm.

### Tip growing cells deprived of functional myosin XI exhibit lateral swelling and drastically reduced growth rate

Functional myosin XI is required for proper tip growth (Ojangu et al., 2007; Peremyslov et al., 2008; Vidali et al., 2010; Park and Nebenfuhr, 2013; Madison et al., 2015) and fluctuation cross-correlation analysis and *in vivo* imaging showed that VAMP-labeled vesicles are a myosin XI cargo (Furt et al., 2013; Bibeau et al., 2020). Nevertheless, the essential mechanism behind myosin XI’s function in polarized cell growth remains to be defined. By exploiting the proper polarized development of myoXIaTS plants at 20°C, we investigated how growth is affected at the single-cell level by the sudden loss of function of myosin XI, upon switching the temperature to 32°C. MyoXIaWT and myoXIaTS grown at 20°C were transferred into an enclosed microscope pre-heated at 32°C and imaged for 5 hours (Movie S1 and MovieS2). The myoXIaTS cells, after exposure to 32°C, exhibit morphological defects as well as growth defects (Figure 4a).

**Figure 4.**
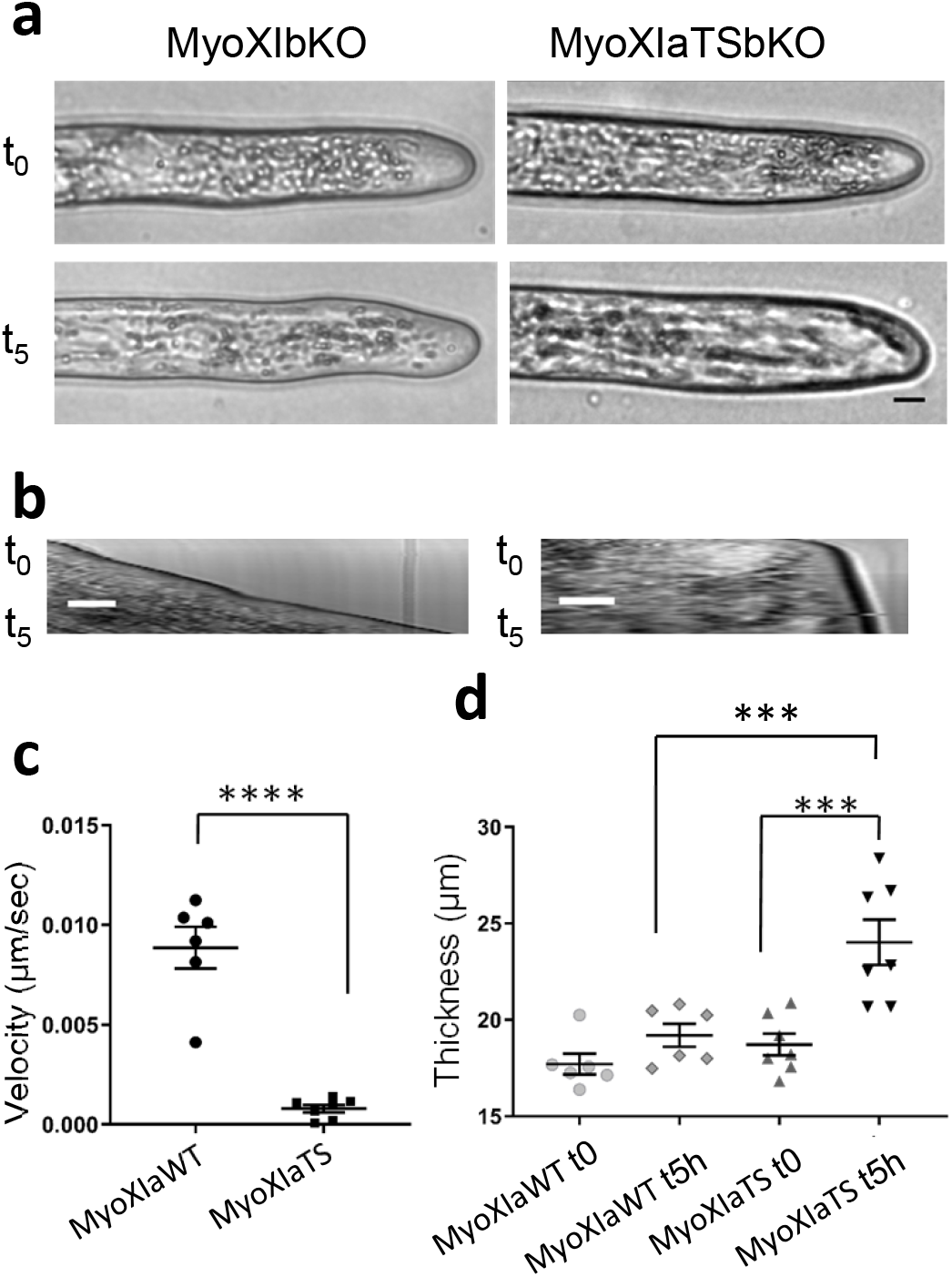
Time-lapse of control and myosin XI TS cells at 32°C show growth and morphology defects of myosin deficient cells. **a.** Representative control and myosin XIaTS cell imaged with epifluorescence microscope enclosed in a temperature-controlled chamber. Cells are imaged for 5 hrs at 32°C. Scale bar = 20 μm. **b.** Representative kymographs depicting cell growth. Scale bar = 10 μm. **c.** Velocity (μm/sec) of control, and myosin XIaTS the cell as quantified from kymographs. Number of cells analyzed: control cells, 6. Myosin XIaTS cells, 7 cells. Difference tested via t-est (*P*< 0.0001), error bars represent SEM. d. Thickness of control and myosin XIa TS cells 50 μm from the cell tip. Number of cells analyzed: control cells, 6. Myosin XIaTS cells, 7 cells. Difference tested via 2 way ANOVA (adjusted *P*= 0.0001), error bars represent SEM.

To quantify changes in the growth rate between the two lines, we performed a kymograph analysis (Figure 4 b,c). MyoXIaTS cells have a drastically reduced growth rate compared to the control. We measured an average growth rate of 8.86 ± 1.04 nm/sec in the myoXIaWT and an approximate ten-fold reduction in growth, to 0.79 ± 0.19 nm/sec, in the myoXIaTS. Visually, the overall morphology of the myoXIaWT cells exposed to 32°C did not change considerably (Figure 4a). The diameter of MyoXIaWT cells only increases modestly, from 17.71 ± 0.54 ◻m (SEM) prior to the incubation, to 19.19 ± 0.6 μm after 5 hours at 32°C,but this change is not significant (Figure 4d). On the other hand, myoXIaTS cells grown at 32°C for 5 hours exhibit considerable lateral swelling in the sub-apex of the cell (Figure 4a,d). The average cell diameter in myoXIaTS cells increases from 18.72 ± 0.56 ◻m prior to the incubation, to 23.02 ± 1.17 ◻m after 5 hr at 32°C, and this change is statistically significant (adjusted *P*<0.01). These observations confirm that myosin XI is critical for polarized growth, and the observed lateral swelling suggests depolarized secretion or alterations in cell wall mechanical properties or both.

### Myosin XIa TS cells exposed at 32°C display higher cell tip curvature and diameter than F-actin depleted cells

To get insights into the individual function of myosin XI and F-actin in maintaining proper cell shape, we investigated how their loss-of-function affects cell morphology. To do so, we performed a time-course fluorescent study (1.5 hrs, 3 hrs, 5 hrs, 8 hrs) in which we treated plants with latrunculin B (20 μM) to depolymerize F-actin or exposed them at 32°C to reduce functional myosin XI. To elucidate myosin XI and F-actin interplay, we then compared the individual myosin XI and the F-actin loss of function phenotypes, with the phenotype of cells deprived simultaneously of both, F-actin and myosin XI. Preliminary analysis showed that, while a phenotype could not be detected at 1.5 hrs, morphological changes manifested at 3 hrs and became more pronounced at 5 hrs and 8 hrs. For this reason, we focused our analysis on the 5 hrs and 8 hrs time points. When plants lack F-actin, the cell tip appears rounder compared to the wild type, predominantly after 5 hrs and 8 hrs of treatment (Figure 5a). Decreased functional myosin XI results in considerable sub-apical swelling and a more narrow tip (Figure 5a), consistent with very slow elongation (Figure 4b).

**Figure 5.**
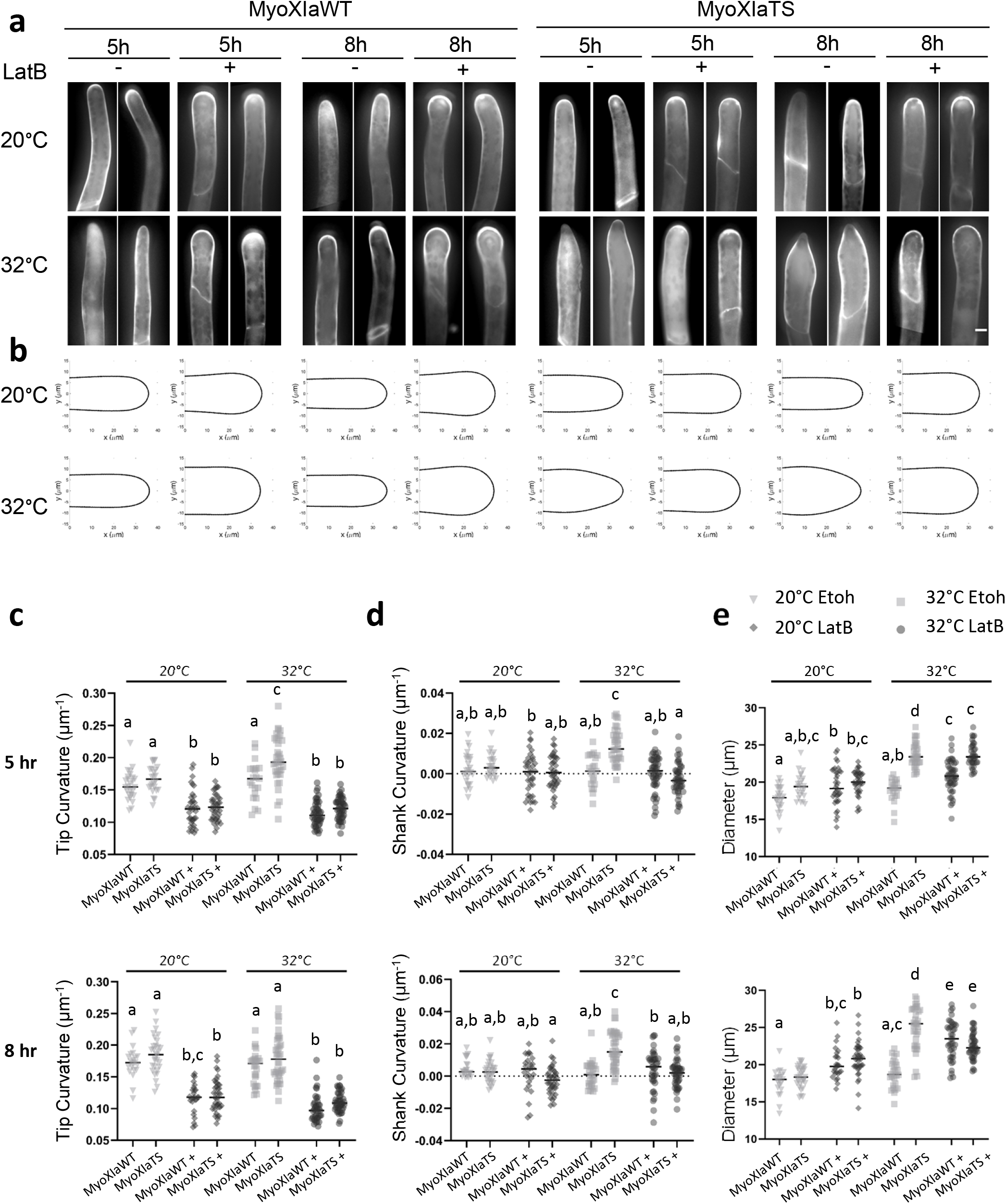
Lack of functional myosin or F-actin affects caulonemata tip morphology differently. **a.** Two representative caulonemata myoXIaWT and myoXIaTS cells grown at 20° C and 32° C with Ethanol (vehicle control) or Latrunculin B (20 μM) before treatment (t0) and for 5 and 8 hours. In the panel, the cellulose is stained with calcofluor-white. Scale bar = 10 μm. **b.** Regeneration of cell contour from computed curvature values. Number of cells used to regenerate each contour: 5 hours, myoXIaWT EtOH 20C, 26. myoXIaWT EtOH 32C, 21. myoXIaWT LatB 20C, 36. myoXIaWT LatB 32C, 47. myoXIaTS EtOH 20C, 24. myoXIaTS EtOH 32C, 35. myoXIaTS LatB 20C, 38. myoXIaTS LatB 32C, 45; 8 hours, myoXIaWT EtOH 20C, 21. myoXIaWT EtOH 32C, 26. myoXIaWT LatB 20C, 26. myoXIaWT LatB 32C, 39. myoXIaTS EtOH 20C, 30. myoXIaTS EtOH 32C, 38. myoXIaTS LatB 20C, 39. myoXIaTS LatB 32C, 43. **c.** average curvature (1/μ) of the cell tip after 5 hrs (top) 8 hrs (bottom). The tip was set at 0 μm, and the curvature between −3 μm to +3 μm from the tip was averaged. **d.** Average curvature (1/μ) of a region on the right side of the cell shank after 5 hrs (top) 8 hrs (bottom). The curvature between 25 μm to 35 μm from the tip was averaged. **e.** Diameter of the cell computed at the widest point after 5 hrs (top) 8 hrs (bottom). Number of cells used for c,d,e, at 5 hrs: myoXIaWT EtOH 20C, 26. myoXIaWT EtOH 32C, 21. myoXIaWT LatB 20C, 36. myoXIaWT LatB 32C, 47. myoXIaTS EtOH 20C, 24. myoXIaTS EtOH 32C, 35. myoXIaTS LatB 20C, 38. myoXIaTS LatB 32C, 45. Number of cells used for c,d,e, at 8 hrs: myoXIaWT EtOH 20C, 21. myoXIaWT EtOH 32C, 26. myoXIaWT LatB 20C, 26. myoXIaWT LatB 32C, 39. myoXIaTS EtOH 20C, 30. myoXIaTS EtOH 32C, 38. myoXIaTS LatB 20C, 39. myoXIaTS LatB 32C, 43. Bars that do not share similar letters denote statistical significance, adjusted *P*<0.05 three-way ANOVA. All values are means ± SEM.

To quantify the observed difference in tip shape, we measured the curvature of the cell at the tip in both caulonemal cells (Figure 5c) and chloronemal cells (Supplemental figure 1c). Following 5 hrs of treatment, the myoXIaTS caulonemal cells at 32°C have higher curvature values than the myoXIaTS cells at 20°C and the myoXIaWT at both temperatures (Figure 5a). In the myoXIaTS, the average tip curvature is 0.16 μm^−1^ ± 0.004 μm^−1^ (SEM) when caulonemal cells are grown at 20°C, and it increases to 0.19 μm^−1^ ± 0.006 μm^−1^ after 5 hrs of exposure at 32°C (Figure 5c, adjusted *P*< 0.01). This curvature difference was only transitory because, after 8 hrs at 32°C, the tip curvature of myoXIaTS cells decreases to 0.18 μm^−1^ ± 0.006 μm^−1^, likely due to the local bulging at the very tip of the cell. MyoXIaTS caulonemal cells treated with latrunculin B for 5 hrs have curvature values of 0.12 μm^−1^ ± 0.003 μm^−1^ both at 20°C and 32°C (Figure 5c). This value is lower than the same cells with unaltered F-actin levels, and the same effect is observed in myoXIaWT cells treated with latrunculin B (Figure 5c). This decrease in curvature is likely the result of a defect in the proper maintenance of anisotropic growth, which causes the rounding of the tip. Similar, but less dramatic effects were observed in chloronemata cells (Supplemental figure 1c).

We also quantified the changes of curvature in the shank, which were visible in the myoXIaTS cells at the restrictive temperature (Figure 5d and Supplemental figure 1d). In these cells, the average curvature on the right side of the shank (similar results were observed on the left side) is 0.014μm^−1^ ± 0.001 μm^−1^ after 5hrs and 0.016μm^−1^ ± 0.002 μm^−1^ after 8hrs (Figure 5d, adjusted *P*< 0.01). On the other hand, in myoXIaTS cells at 20°C and myoXIaWT cells at both temperatures, the shank does maintain a straight morphology (~ zero curvature).

Depolymerization of F-actin by latrunculin B treatment in both TS and WT cells does no result in an increase of lateral curvature at any temperature, suggesting that F-actin is necessary for the myosin XI loss-of-function phenotypes observed. Again, similar, but less dramatic effects were observed in chloronemata cells (Supplemental figure 1d).

This assay also allowed us to confirm the increase in cell thickness following myosin XI loss-of-function (Figure 2a,d). In myoXIaTS caulonema cells, cell thickness increases from 19.45μm ± 0.34 μm^−1^ at 20°C, to 23.66μm ± 0.26 μm^−1^at after 5 hrs at 32°C, and to 25.33 μm ± 0.520 μm^−1^ after 8 hrs at 32°C (Figure 5e, adjusted *P*< 0.01), while the myoXIaWT cells do not exhibit a significant diameter change from 20°C to 32°C (Figure 5e). Only a modest change in thickness was observed after latrunculin B treatment at high temperature, but the magnitude of the changes was smaller than that produced by reducing myosin XI function in the presence of F-actin. Together, our results show that myosin XI and F-actin play separable and sometimes complementary roles in cell growth and morphogenesis.

### Lack of functional myosin XI dissipates tip-localized VAMP-labeled vesicles and F-actin within minutes

*P. patens* apical cells exhibit an enrichment of myosin XI and F-actin at the apex, which is both essential for tip growth and only present in cells that are actively growing (Vidali et al., 2009a; Vidali et al., 2010). Similarly, a population of VAMP-labeled vesicles populates the tip (Furt et al., 2013; Bibeau et al., 2018; Bibeau et al., 2020), while myosin XI and VAMP-labeled vesicles accumulation lead the F-actin’s accumulation by 18.6 sec at the tip of the cell (Furt et al., 2013). This prompted us to hypothesize that myosin XI functions by clustering vesicles at the cell’s tip, and that this clustering is important for F-actin organization and vesicle accumulation, both necessary for growth (Furt et al., 2013; Bibeau et al., 2018). To evaluate this hypothesis and to investigate the cause for the drastic reduction in the observed growth rate of myoXIaTS cells at 32°C, we explored the dependence of the apical localization of VAMP-labeled vesicles and F-actin on functional myosin XI. To do this, we transformed the myoXIaWT and myoXIaTS lines with the vesicle marker 3xmCherry-VAMP72A1 (referred to as myoXIaWT-VAMP and myoXIaTS-VAMP lines); these lines also express the F-actin probe, Lifeact-mEGFP (Vidali et al., 2009a). As expected, at 20°C and 32°C, myoXIaWT cells exhibit the characteristic VAMP-labeled vesicle and F-actin accumulations at the cell tip (Figure 6). However, in the myoXIaTS cells exposed 32°C for only 20 minutes, the vesicle cluster dissipates, and concomitantly, the F-actin spot disappears. These results are consistent with myosin XI functioning in the clustering of vesicles at the cell tip, and with this clustering being important to maintaining F-actin organization and polarized secretion needed for tip growth.

**Figure 6.**
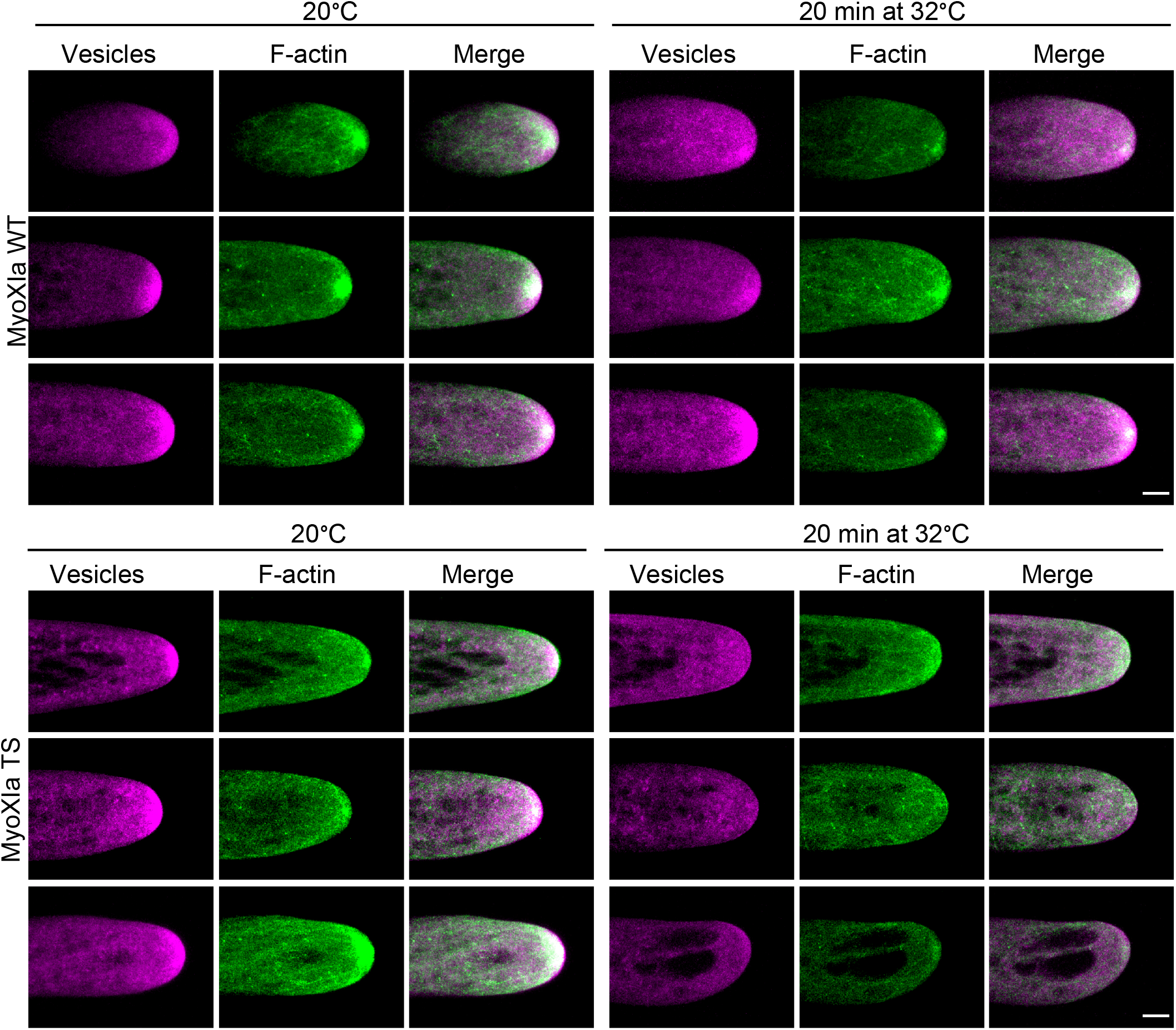
F-actin spot and tip oriented vesicle cluster dissipate upon 20 minutes exposure at 30°C. Myosin XIbKO 3xmCherry-VAMP71A1 and myosinXIa TS 3xmCherry-VAMP72A1 imaged while exposed at 20 °C and 32 °C. In the panel, green represents LifeAct-GFP labeled F-actin and magenta the 3xmCherry-VAMP72A1 secretory vesicles. MyoXIaTSbKO cells exposed at 32 °C for 20 minutes fail to exhibit vesicle clustering at cell tip and F-actin spot. For display purposes, the images were enhanced by contrast normalization (0% saturation), gaussian blur (0.6%), and manual adjustment of brightness and contrast levels. Scale bar 5 μm.

### Myosin XI is essential for the formation of latrunculin B-induced vesicle clusters

In previous work, Furt et al. observed that upon treatment with very low concentrations of latrunculin B, myosin XI clusters followed by F-actin polymerization arise in ectopic sites at the cell’s shank, recapitulating myosin XI and F-actin correlation observed at the growing tip (Furt et al., 2013). Based on these published observations, and on our aforementioned fluorescent results (Figure 6), we hypothesize these ectopic clusters are the result of myosin XI’s role in clustering vesicles and that this clustering results in F-actin local polymerization, and subsequent actin-polymerization dependent vesicle motility. To test our hypothesis, we used the myoXIaWT-VAMP and myoXIaTS-VAMP lines and attempted to reproduce the previously observed myosin XI and F-actin ectopic clusters upon latrunculin B treatment (Furt et al., 2013), both at the permissive and restrictive temperature. Importantly, the use of a myosin XI temperature-sensitive allele allowed us to test if myosin XI is necessary for the observed ectopic vesicle clustering.

When treating the myoXIaWT-VAMP and myoXIaTS-VAMP with latrunculin B (0.25 μM) at 20°C, we observed, in multiple locations in the cell, an enrichment of vesicle-clusters followed by F-actin polymerization (Figure 7). When F-actin is polymerized, the VAMP-labelled vesicles clusters become motile and propel in the cell, consistent with an F-actin polymerization dependent motility. When we treated myoXIaTS-VAMP cells with latrunculin B (0.25 μM) and exposed them at 32°C, the VAMP-labeled vesicle clusters fail to form. Importantly, the control myoXIaWT-VAMP cells treated with latrunculin B and exposed at 32°C still exhibit the motile vesicle clusters (Figure 7 and Movie S3 and S4). In myoXIaWT cells, 12 out of 14 cells at 20°C and 15 out of 15 cells at 32°C exhibited clusters and the difference is not significant (Fisher’s exact test, two-tailed *P*= 1); in myoXIaTS cells, 13 out of the 15 cells at 20°C exhibited clusters, while only 1 out of 15 cells exhibited clusters at 32°C and the difference is significant (Fisher’s exact test, two-tailed *P*<0.0001) (Figure 7 and Movie S5 and S6). These results prove that myosin XI is essential for the formation of ectopic vesicles enrichments, hence to cluster VAMP-labeled vesicles resulting in the subsequent polymerization of F-actin. Based on the observed similarities between the apical vesicle cluster and the ectopic clusters, including their dependence on myosin XI for their formation, we propose that a similar myosin XI-dependent vesicle-clustering mechanism is responsible for both.

**Figure 7.**
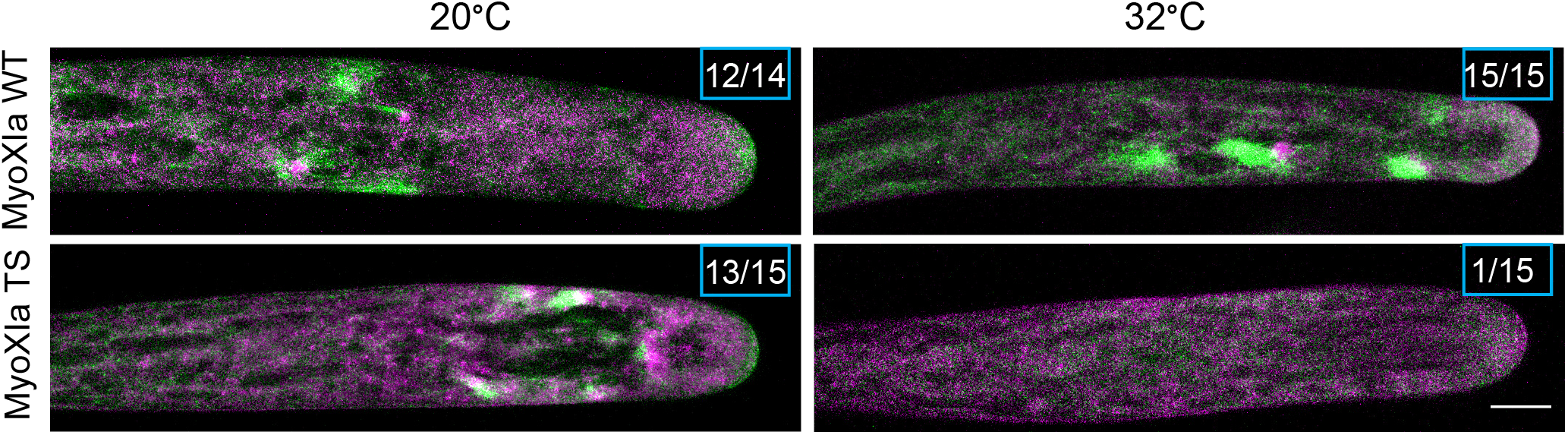
Low concentration Latrunculin B-induced motile vesicles clusters followed by F-actin are myosin XI dependent. Representative myosin XIbKO 3xmCherry-VAMP71A1 and myosin XIa TS 3xmCherry-VAMP72A1 cells treated with 0.25 μM LatB and imaged while exposed at 20 °C and 32 °C. In the panel, green represents LifeAct-GFP labeled F-actin and magenta the 3xmCherry-VAMP72A1 vesicles. For display purposes, the images were enhanced by γ-adjustment (1.3%), gaussian blur (0.5%), and manual adjustment of brightness and contrast levels. Number of cells analyzed: control 20 °C, 14. Control 32 °C, 15. TS 20 °C, 15. TS 32 °C, 15. In the top right corner, it is reported the number of cells exhibiting the ectopic clustered compared to the total number of cells analyzed. The difference in the numbers of observed vesicle clusters is not significant between the two cell lines at 20 °C. The difference in numbers of observed vesicle clusters is highly significant between the two cell lines at 32°C (Fisher’s exact test, two-tailed *P*<0.0001). Scale bar 10 μm.

## DISCUSSION

Here we have revealed essential functions of myosin XI in plant cells. We showed that myosin XI is critical for polarized growth via its vesicle clustering activity, which induces actin polymerization. We also showed that this myosin is important to maintain cell morphology, vacuole structure, and cell survival. These conclusions are supported by experimental evidence obtained using a temperature-sensitive allele of myosin XI that we generated. The conclusion that myosin XI participates in vesicle clustering, and that this clustering is important for actin polymerization, is consistent with the hypothesis that vesicle clustering plays an essential role in protonemal tip growth. It will be interesting to investigate if a similar vesicle-clustering function and vesicle-mediated actin polymerization process exist in other plant tip growing cells. The use of a myosin XI TS allele provided an exceptionally advantageous tool to overcome growth limitations resulting from transient RNAi (Vidali et al., 2010). Using this tool, we were able to analyze phenotypes of differentiated cells, specifically caulonemata, that do not develop in plants undergoing RNAi of myosin XI.

In line with the hypothesis, we showed myosin XI is essential for vesicle clustering and F-actin accumulation at the cell tip. This result provides new insight into the relationship between myosin XI, vesicles, and F-actin in maintaining persistent polarized growth in plants. Our data indicate that myosin XI is the driver of a long-range F-actin polymerization-driven vesicular transport system, in which myosin XI binds secretory vesicles that carry an F-actin nucleator, such as class II formins (Vidali et al., 2009c) or the ARP2/3 complex (Perroud and Quatrano, 2006) or both. Subsequent polymerization of F-actin serves as a binding route for more motors, in a myosin XI-driven feedback that locally increases F-actin. Our data fit with existing evidence in *P. patens*, in which motile structures propelled by F-actin originating from formin II clusters have been described (van Gisbergen et al., 2012; Wu and Bezanilla, 2018). In addition, in support of our model, formin II localizes at the apex of tip growing cells, and it is essential for polarized growth (Vidali et al., 2009c). A similar vesicular transport has been described in mice oocytes (Schuh, 2011), and vesicle clustering and actin-based propulsion have been previously observed in other plants; for instance, in *A. thaliana* pollen tubes, vesicle clusters propelled by formin-generated F-actin is essential for germination (Liu et al., 2018). However, inhibition of myosin-F-actin interaction via BDM suggested the mechanism observed by Liu et al. is myosin XI-independent (Liu et al., 2018), indicating that a fundamental difference may exist in the vesicle clustering and transport between the moss *P. patens* protonemata and the pollen of the vascular plant *A. thaliana*. On the other hand, it was recently shown that a myosin XI from the bryophyte *Marchantia polymorpha* could rescue many of the developmental phenotypes associated with the loss of function of myosin XI in *A. thaliana* (Duan et al., 2020). This results suggest a high degree of functional conservation between non-vascular and vascular plant myosin XI proteins.

Our study demonstrates a link between myosin XI function and polarized cell’s growth-rate and morphology. Myosin XI depleted cells exhibit a highly reduced growth rate, consistent with diffusion-based growth (Bibeau et al., 2018), and a simultaneous increase in cell thickness. The reduced growth-rate could be due to a small amount of myosin XI retaining its function. Furthermore, our results show that depletion of myosin XI results in pointier cells with higher tip curvature and F-actin depleted cells result in blunter cells with lower tip curvature values. These results directly relate a molecular motor and the cytoskeleton to cell shape and provide us with a powerful tool to further study the regulation of cell morphology. Mathematical modeling of tip growing cells identified scaling laws that connect cell wall geometry to internal parameters (Campàs and Mahadevan, 2009; Campas et al.). Specifically, Campas et al. described a scaling law that relates the cell radius and the apical radius of curvature to the size of the vesicle secretion area (Campas et al., 2012). According to Campas et al., in blunt cells, the cell radius is proportional to the radius of curvature at the tip, and this implies that the secretion area scales with cell size; hence, it increases with an increase of cell size. On the other hand, in pointy cells, the radius of tip curvature is constant and independent of the cell’s radius, and this implies the secretion area is constant (Campas et al., 2012). Integrating this with our data, we speculate that in myosin XI TS cells, secretion is highly reduced, but confined to a constant area in the cell’s apex (Bibeau et al., 2018), resulting in a highly reduced growth rate, very thin cell diameter and a high tip curvature. On the other hand, in F-actin depleted cells, vesicles are passively transported via diffusion. Untargeted vesicle diffusion results in docking in a broader area at the tip, causing growth arrest and lateral growth and increase of cell thickness.

Alongside with deepening our understanding of myosin XI function in cell growth, we showed myosin XI is important for vacuole structure and cell viability. Furthermore, our data indicate depolymerization of F-actin attenuates the myosin XI-dependent cytological aberrations, opening new lines of investigation for the myosin XI-F-actin system in tip growing plant cells. The evidence that the F-actin myosin system could regulate vacuole structure in *P. patens* is inconsistent with previous reports. In fact, Oda et al. showed that microtubules are responsible for vacuole structure regulation in *P. patens* (Oda et al., 2009). However, the investigation of Oda et al. is limited to chloronemata and rhizoid cells, while our focus is on caulonemata cells. Caulonemata exhibit a lower density of organelles, especially chloroplasts, and have a higher growth rate than chloronemata (Furt et al., 2012), and this could result in a different vacuolar regulation system. Despite differences with previous reports in *P. patens*, our observations are common to the flowering plants *A. thaliana* and *N. tabacum*, in which the F-actin cytoskeleton is responsible for regulating vacuolar structure (Higaki et al., 2006; Scheuring et al., 2016). Furthermore, an F-actin-myosin V system regulates vacuole inheritance in *S. cerevisiae* (Tang et al., 2003). Further experiments are needed to broaden our understanding of myosin XI-F-actin involvement in vacuole regulation in plants. Identification of myosin XI interactors on the tonoplast membrane could confirm myosin XI direct role in vacuole transport in the cell. The heavy misregulation of the vacuole after 24 hrs of myosin XI depletion could be one of the causes of loss of cell viability. The lack of myosin XI has been previously related to premature cell senescence and cell death in *A. thaliana* (Ojangu et al., 2018). In discordance with our results, previous studies link F-actin depolymerization with cell death. In fact, in *A. thaliana*, latrunculin B induced F-actin depolymerization causes programmed cell death in pollen tubes (Thomas et al., 2006). Furthermore, Ojangu et al. show disruption of F-actin via latrunculin B results in increased transcriptional levels of stress-related genes (Ojangu et al., 2018). In the yeast, *S. cerevisiae*, decreased F-actin dynamics result in cell death, while the increase in F-actin dynamics increases in lifespan by 65% (Gourlay et al., 2004). Further research is needed to clarify why the depolymerization of F-actin rescues cell viability in *P. patens*. This study demonstrates a novel role for myosin XI in vesicle clustering and finds an association between myosin XI function with the maintenance of vacuole structure and cell viability.

## METHODS

### Generation of a *P. patens* myosin XI temperature-sensitive allele and a myosin XIb KO line

To identify residues that render myosin XI temperature-sensitive, we first introduced single mutations in the hydrophobic core of the myosin XI motor domain via PCR site-directed mutagenesis (Vidali et al., 2009b). Three different hydrophobic residues (L79, V584, L616) were initially mutagenized, and three different constructs were tested. The mutagenesis was performed on the entry clone pENT-MyoXIaCDS harboring the ORF of the myosinXIa cDNA (Vidali et al., 2010). Three PCR reactions were performed using a forward mutagenic primer and the reverse primer M13 in the three following primer combinations: MyoXIA-L616A-F and M13R, MyoXIA-V584A-F and M13R, MyoXIA-L79A-F, M13R (see supplementary table). Three additional PCR reactions were performed using a reverse (non-mutagenic) primer and the forward primer M13 in the three following combinations: MyoXIA-L616A-R and M13F, MyoXIA-V584A-R and M13F, MyoXIA-L79A-R, M13F. The PCR fragments and pENT-MyoXIaCDS were digested with NotI and AscI and ligated together, generating three different constructs: pENTMyoXIaL79A, pENTMyoXIaV584A, pENTMyoXIaL616A. The ligated constructs were verified by sequencing. An LR Gateway reaction was used to transfer the pENTMyoXIaL79A, pENTMyoXIaV584A, and pENTMyoXIaL616A to the expression vector pTHUbiGate and generated three expression clones. The expression clones were co-transformed in moss with an RNAi construct targeting both myosin XIa, and XIb from their 5’UTRs (Vidali et al., 2010) and their ability to complement at 20-25°C and plant temperature sensitivity at 32°C was tested.

The constructs harboring MyoXIaV584A and MyoXIaL616A complemented at 25°C and showed mild temperature sensitivity at 32°C. The pENTMyoXIaL79A construct did not show any temperature sensitivity and was not pursued any further. To obtain a stronger temperature sensitivity, we generated a construct harboring both V584A and L616A. In order to do so, we used a cloning strategy taking advantage of the presence of a MfeI restriction site between the two sites that need to be mutated. We digested pENTMyoXIaV584A with KpnI and MfeI and kept the smaller fragment, which contained the V584A substitution; we cut pENTMyoXIaL616A with KpnI and MFeI and kept the large fragment, which contained the vector that also had the L616A mutation. The two fragments were ligated together (pENTMyoXIaV584AL616A). An LR Gateway reaction was used to transfer the construct from pENTMyoXIaV584AL616A to pTHUbiGate. The expression clone was tested via complementation assay, and preliminary analyses showed complementation at 20°C and significant temperature sensitivity at 32°C.

To introduce the mutations in the myosin XIa locus, a knock-in construct was generated by separately cloning the ~1 kb regions 5^’^ and 3^’^ to the section to be mutagenized. These regions were then assembled with the sequences from the ORF clone pENTMyoXIaV584AL616A. For the 5’ homologous arm, a forward primer (MyoXIAKI1F) hybridized on Intron 7 and generated a PCR product with a CACC TOPO-cloning site followed by a SwaI restriction site, a reverse primer (MyoXIAKI1R) hybridized on exon 15 introducing a SalI restriction site through the introduction of a silent mutation. This PCR product was cloned into a TOPO cloning vector (pENTR/D-TOPO) (myoXIa5^’^arm-TOPO). For the 3’ homologous arm, the forward primer (MyoXIAKI2F) hybridized on exon 15 and generated a PCR product with a CACC TOPO-cloning site followed by a SalI restriction site, and the reverse primer (MyoXIAKI2R) hybridized on exon 22 and introduced a SwaI restriction site. This PCR product was cloned into a TOPO cloning vector (myoXIa3^’^arm-TOPO). The third construct was generated by performing PCR on pENTMyoXIaV584AL616A. The forward primer (MyoXIAKI2F) hybridized on exon 15 and generated a SalI restriction site, and the reverse primer (MyoXIAEX18R) hybridized on exon 18, downstream an XhoI site. This PCR product was cloned into a TOPO cloning vector (V584AL616A-TOPO), and it contains the two mutations V584A and L616A (respectively on exon 16 and 17), and it does not contain introns 15, 16 and 17. The 3^’^ arm-TOPO and V584AL616A-TOPO vectors were digested with SalI/XhoI; the small fragment of the myoXIa3^’^ arm-TOPO and the larger fragment of the V584AL616A TOPO were ligated. The resulting plasmid and the myoXIa5’ arm-TOPO vector were digested with SalI/AscI. The smaller fragment of the first digestion was ligated with the larger fragment of the myoXIa5’ arm-TOPO. The ligations generated the final temperature-sensitive knock-in construct (pTOPO-MyoXIA-TSKI). Since, in *P. patens*, myosin XIa and myosin XIb are functionally redundant (Vidali et al., 2010) together with pTOPO-MyoXIA-TSKI we co-transformed a myosin XIb knock-out construct (L1L2-MyoXIB-KO) (Sun et al., 2018), which, by homologous recombination, replaces the myosin XIb gene with a hygromycin cassette. Both constructs were linearized by SwaI digestion and transformed into protoplasts of a moss line expressing Lifeact-mEGFP (Vidali et al., 2009a). Afterward, plants were regenerated as previously described (Liu and Vidali, 2011).

To select plants in which both the knock-in and knock-out homologous recombination reactions happened successfully, we used a PCR genotyping strategy. From single clones, we extracted the genomic DNA (Quick-DNA Plant/Seed Miniprep Kit, Zymoresearch) and performed PCR. To verify the presence of the myosinXIaTS allele, we used a forward primer (MyoXIAIntron6F) that hybridized on Intron 6, and the reverse primer (MyoXIAIntron18R) which hybridized on Intron 18. These primers amplify a 3.2 Kb band in plants with the myosin XIaTS allele and a 3.7 Kb band in the plants with the wild type allele. To simplify the discrimination between these two bands, we took advantage of the presence of a SalI site in the knock-in construct and digested the PCR product with SalI. After digestion, the 3.2Kb band was cut into two bands of 2.7 Kb and 0.5 Kb. To verify the insertion of the hygromycin cassette in the myosin XIb gene, a forward primer (MyoXIbEXTGnmc5F) hybridized upstream to exon 1 and a reverse primer (NosLoxR) hybridized on the Lox site on the inserted construct (Sun et al.). These primers amplify a 1.9 Kb band only in plants in which the hygromycin cassette has been inserted in the myosin XIb gene. All reported moss lines are available upon request.

### Plant morphometric assay

*P. patens* protoplasts of the myoXIaWT and myoXIaTS lines were regenerated for four days at 25°C (permissive temperature) in PpNH_4_ medium containing 10 mM CaCl_2_ and 6% Mannitol. After regeneration, protoplasts were transferred to plates containing growing medium (PpNH_4_) and placed in a 32°C chamber or 20°C chamber for three days. All growth chambers were set with a 16 hrs light 8 hrs dark cycle. Plants were stained with calcofluor-white (Fluorescent Brightener 28, Sigma) at a final concentration of 10 μg/ml, to mark their outline and imaged with an inverted epifluorescence microscope Axiovert 200 M (Zeiss) and a 10 X lens (0.25 NA). Imaging was performed with the AxioVision (Zeiss) software. The calcofluor-white signal was imaged with a DAPI filter. Plant area and solidity were measured with an ImageJ Morphology macro (Galotto et al., 2019).

### Time-course fluorescent image acquisition

*P. patens* protoplasts of myoXIaWT and myoXIaTS lines were regenerated for three days at 25°C in PpNH_4_ medium containing 10 mM CaCl_2_ and 6% Mannitol covered with cellophane. The plants were then transferred on PpNH_4_ moss medium at 20°C for 4 days, then transferred on PpNO_3_ medium for additional 4 days at 20°C. This procedure allows for the homogenous development of single moss plants and the generation of tissue rich in caulonema cells. Plants were then transferred on control media, or on media containing latrunculin B (Sigma) at a final concentration of 20 μM. Plants were then exposed at 20°C or 32°C for 1.5 hrs, 3 hrs, 5 hrs, 8 hrs. At each time point, plants were transferred on a microscope slide and mounted on an agar pad for imaging. To image the outline of the plant, 30 μl of calcofluor-white (final concentration of 10 μg/ml) was pipetted on the plants. Plants were imaged with an inverted epifluorescence microscope Axiovert 200 M (Zeiss). The calcofluor-white signal was imaged with a DAPI filter. To detect a high-quality image of the plants, we used an automated technique recently developed in our lab (Galotto et al., 2019). Briefly, we used the MosaiX AxioVision software in combination with an automatic stage that allowed us to save the position of different plants in the microscope slide. For each plant, we imaged a stack of 5 planes 20 μm apart. Images were then processed by an ImageJ macro we developed (available upon request). The macro stitches together the acquired tiles by using an imageJ plugin called “Grid/Collection stitching plugin” (Preibisch et al., 2009) and performs the maximum projection of the plants.

### Detection and analysis of cell tip curvature

To obtain a metric for quantification of the cell tip shape, we used the imageJ plugin “J Filament” (Smith et al., 2010). By using an in-house ImageJ macro to crop and rotate tips (available upon request), we first cropped single tip cells from the micrographs generated as described in the “Image acquisition” section. All the cells were rotated such that the very tip section of the cells point perpendicularly to the right. We inverted the color of the cropped images in ImageJ and used “Find Edges” command to highlight cell contours. We then proceeded to trace the contours using the JFilament plugin, which gives as an output the {*x,y*} coordinates along the outline. The {*x,y*} coordinates then underwent a linear interpolation using a custom MATLAB code to obtain coordinates with equal spacing of 1 pixel (0.489 μm). To quantify the cell morphology, we developed a MATLAB code (available upon request), which takes the {*x,y*} coordinates as input and measures the curvatures along the cell peripheries as a function of the distance from the tip. To avoid inherent noise from experiments and noise introduced by image processing, instead of calculating the curvature of each coordinate triplet from the two bonds connecting the coordinates to its adjacent neighbor, we calculated the curvature using longer bonds. To achieve this, we first drew lines connecting all coordinates. We then drew a circle of radius Δs centered at each of these coordinates. Two new bonds were constructed for each coordinate by drawing lines from the coordinate point (center) to the points where the circle intersects with the contour lines. This process is illustrated in Fig. S2. We approximated curvature, *K*(*S*), as the angle change with respect to distance, given as

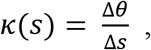

where Δθ is the change in angle between the two bonds. By doing a parameter scan, we decided that Δs of 4 μm are optimal for noise elimination while still retaining information from cell contours (Fig. S3). In addition, each cell was scanned perpendicularly along the x-axis up to contour length of 40 μm from the tip, to find the largest diameter (Fig S4).

### Bright-field time-lapse imaging of cells transferred to high temperature

One week old myoXIaWT and myoXIaTSplants, cultured at 20°C, were transferred to a 35 mm glass-bottom dish (Matsunami) suitable for imaging. The dish contains a thick outer layer of agar and a thin inner layer of agar on top of which the plant is positioned. After the plants are transferred, the chambers are placed in the 20°C incubator for 1 week to let the moss adapt, prior to imaging. To image tip growing cells while growing at 32°C, we built a microscope enclosure equipped with a heating system. Bright-field time laps were acquired via a Zeiss Axio Observer.A1, with a 20X objective lens (N.A. 0.3). Images are acquired with the software Micromanager (Edelstein et al., 2014), every 5 minutes for 5 hours. To measure cell growth we generated a kymograph for each cell with the imageJ “multi kymograph” plugin.

### Confocal imaging

To observe F-actin and secretory vesicles in the TS background, the myoXIaWT and myoXIaTS protoplasts were transformed with the pTK-Ubi-3xmCherry-VAMP, which drives protein expression from a constitutive promoter and targets a redundant copy of the ARPC2 gene (Furt et al., 2013). To induce the formation of the ectopic F-actin vesicle clusters (Furt et al., 2013), single myoXIaWT-VAMP72A1 and myoXIaTS-VAMP72A1 plants were placed in a round 35 mm confocal dishes (VWR), on top of an agar pad containing latrunculin B (0.25 μM); 20 ul of liquid moss medium containing latrunculin B (0.25 μM) was pipetted on the plants and a small round coverslip secured with VALOP was used to close the preparation. Clusters were imaged with the Leica SP5 scanning confocal microscope. To image cells while exposed to 20°C, the petri dish was placed in a holder connected to a temperature control system (Warner Instruments CL-200) set to 20°C. To image cells while exposed at 32°C, the temperature control system was set to 32°C and a lens heater (Warner Instruments) was placed on the oil immersion 63X lens (N.A. = 1.4) used for imaging. Prior to imaging, dishes were pre-exposed to 32°C for 15 minutes in an incubator. The temperature in the preparation was detected via a thermocouple placed in proximity to the plant. The 488 and 561 nm laser were used to excite mEGFP and mCherry respectively. The double dichroic 488/561 was used, with bidirectional X laser scanning and pinhole of 2. Z-stacks of 4 optical slices separated by ~6 μm were acquired at 3 sec intervals at 200 Hz laser scanning speed.

To image F-actin and secretory vesicles at the cell tip, the same microscopy preparations and temperature control systems were used, with the only difference that 2 ml of mineral oil (VWR) were placed on top of the sealed prep to facilitate temperature detection with a thermocouple. Preparations were kept at 20°C until placed in the microscope stage, and the same cell was imaged first at 20°C and then at 32°C. Z-stacks of 10 optical slices were acquired at 200 Hz laser speed, with a pinhole at 1 AU. After acquiring the Z-stack at 20°C, the temperature was switched to 32°C, and the second Z-stack was acquired 20 minutes after the temperature in the chamber reached 32°C. Images were enhanced by contrast normalization (0% saturation), gaussian blur (0.6%), and manual adjustment of brightness and contrast levels in ImageJ for display purposes.

### Vacuolar staining

The vacuole was stained using the marker MDY-64 (ThermoFisher). The stock solution was prepared in DMSO at 250 μM and diluted in liquid plant medium to a working concentration of 500 nM. *P. patens* plants were exposed at 20°C or 32°C for 2 hrs, then incubated for 15 minutes with the dye, which was subsequently removed by washing twice with liquid plant medium. The samples were imaged with a Leica SP5 Point Scanning Confocal Microscope. MDY-64 was excited via the 488 nm argon laser, the cells imaged with a 63X lens (N.A. 1.4), and a double dichroic mirror 458/514. For display purposes, the images were γ-adjusted (0.8%) with ImageJ.

### Cell death assay

To perform this experiment, we used the same settings described in “Image acquisition” section, but prolonging exposure at the stated conditions to 24 hrs. Plants were stained with both calcofluor-white at a final concentration of 10 μg/ml, and propidium iodide (Thermo Fisher Scientific) at a final concentration of 20 μg/ml. The calcofluor-white and propidium iodide signals were detected with the DAPI and rhodamine filters, respectively. The number of dead cells per plant was estimated by counting of propidium iodide stained nuclei.

### Statistical methods

For comparison of multiple treatments, the following statistical tests were performed. A two-way analysis of variance (morphometric assay), a three-way analysis of variance (cell death assay), t-test (bright-field time-lapse imaging of cells transferred to high temperature), and Fisher’s exact test (confocal imaging) were computed with the Software Graphpad Prism version 8.4.2 for Windows (GraphPad Software, La Jolla California USA, www.graphpad.com). A three-way analysis of variance (cell tip curvature and diameter) was performed with R software (www.R-project.org). A comparison of means was done by the Tukey post hoc tests; an adjusted *P*-value of 0.05 was used as significant. Specific information about the number of samples analyzed and the comparisons are specified in the figure legend.

## Acknowledgments

The autors would like to thank all the members of the Vidali Lab and Tüzel Lab for providing precious feedback on the manuscript and Victoria Huntress for her support with the use of the Microscopy core facility at Worcester Polytechnic Institute. In addition, we would like to thank Dr. Callum J. Bell from the National Center for Genome Resources for assistance in the preliminary transcriptome analysis; the collaboration with Dr. Bell was supported by the Gordon and Betty Moore Foundation (grant number 4823). This work was supported by the National Science Foundation (NSF-MCB,1253444 to L.V.).

## Author Contributions

L.V., G.G., J.B.P designed the research. G.G, J.B.P, P.W., Y.C.L. E.T. conducted experiments and analyzed the data. P.J.S. collaborated in analyzing the data. G.G, J.B.P, P.W., E.T. interpreted the data. G.G. wrote the initial draft of the manuscript and L.V., J.B.P edited the draft. All authors reviewed and approved the final version of the manuscript.

---

**Supplemental Figure S1. Lack of functional myosin or F-actin affects chloronemata tip morphology differently. a.** Two representative chloronemata myoXIaWT and myoXIaTS cells grown at 20° C and 32° C with Ethanol (vehicle control) or Latrunculin B (20 μM) before treatment (t0) and for 5 and 8 hours. In the panel, the cellulose is stained with calcofluor-white. Scale bar = 10 μm. **b.** Regeneration of cell contour from computed curvature values. Number of cells used to regenerate each contour: 5 hours, myoXIaWT EtOH 20C, 36. myoXIaWT EtOH 32C, 30. myoXIaWT LatB 20C, 47. myoXIaWT LatB 32C, 37. myoXIaTS EtOH 20C, 31. myoXIaTS EtOH 32C, 47. myoXIaTS LatB 20C, 41. myoXIaTS LatB 32C; 8 hours, myoXIaWT EtOH 20C, 31. myoXIaWT EtOH 32C, 34. myoXIaWT LatB 20C, 35. myoXIaWT LatB 32C, 27. myoXIaTS EtOH 20C, 23. myoXIaTS EtOH 32C, 31. myoXIaTS LatB 20C, 35. myoXIaTS LatB 32C, 35. **c.** Average curvature (1/μ) of the cell tip after 5 hrs (top) 8 hrs (bottom). The tip was set at 0μm, and the curvature between −3 μm to +3 μm from the tip was averaged. **d.** Average curvature (1/μ) of a region on the right side of the cell shank after 5 hrs (top) 8 hrs (bottom). The curvature between 25 μm to 35 μm from the tip was averaged. **e.** Diameter of the cell computed at the widest point after 5 hrs (top) 8 hrs (bottom). Number of cells used for c,d,e, at 5 hrs: myoXIaWT EtOH 20C, 36. myoXIaWT EtOH 32C, 30. myoXIaWT LatB 20C, 47. myoXIaWT LatB 32C, 37. myoXIaTS EtOH 20C, 31. myoXIaTS EtOH 32C, 47. myoXIaTS LatB 20C, 41. myoXIaTS LatB 32C, 38. Number of cells used for c,d,e, at 8 hrs: myoXIaWT EtOH 20C, 31. myoXIaWT EtOH 32C, 34. myoXIaWT LatB 20C, 35. myoXIaWT LatB 32C, 27. myoXIaTS EtOH 20C, 23. myoXIaTS EtOH 32C, 31. myoXIaTS LatB 20C, 35. myoXIaTS LatB 32C, 35. Bars that do not share similar letters denote statistical significance, adjusted *P*<0.05 three-way ANOVA. All values are means ± SEM.

**Supplemental Figure S2. Illustration of the curvature calculation along the cell periphery.** A circle of radius Δs is drawn around each coordinate, where two bonds are constructed from by drawing lines from the coordinate point (center) to the points where the circle intersects with the contour lines. Curvature is calculated as the change in angle between the two bonds over the new bond length, as described in the methods.

**Supplemental Figure S3. Determination of optimal bond spacing for curvature calculations.** Curvatures as a function of contour distance from the tip for 5 sample cells are plotted for different bond spacings. The spacing of 4 μm, which is optimal for noise elimination while still retaining information from cell contours, was selected for all curvature calculations presented in this work.

**Supplemental Figure S4. Illustration of the maximum width determination.** For each cell, we performed horizontal scans from the tip at every 0.5 μm along the x-axis of the cell up to the contour length of 40 μm. Red indicates the scan to the top contour of the cell, and blue indicates the scan to the bottom contour of the cell. The longest length from the top contour to the bottom contour (red bar + blue bar) is reported as the width of the cell.

**Movie S1: Growth rate and cell morphology of a control cell at 32°C are not affected by the temperature.** Representative control cell imaged with an epifluorescence microscope enclosed in a temperature-controlled chamber. Cells are imaged for 5 hrs at 32°C. Scale bar = 20 μm.

**Movie S2: Myosin XI TS cells at 32°C exhibit slow growth and morphology defects.** Representative myosin XI TS cell imaged with epifluorescence microscope enclosed in a temperature-controlled chamber. Cells are imaged for 5 hrs at 32°C. Scale bar = 20 μm.

**Movie S3: Representative myosin XIbKO 3xmCherry-VAMP71A1 cell treated with 0.25 μM LatB and imaged while exposed at 20 °C.** Green represents LifeAct-GFP labeled F-actin and magenta the 3xmCherry-VAMP72A1 vesicles. Scale bar 10 μm.

**Movie S4: Representative myosin XIbKO 3xmCherry-VAMP71A1 cell treated with 0.25 μM LatB and imaged while exposed at 32 °C.** Green represents LifeAct-GFP labeled F-actin and magenta the 3xmCherry-VAMP72A1 vesicles. Scale bar 10 μm.

**Movie S5: Representative myosin XIa TS 3xmCherry-VAMP72A1 cells treated with 0.25 μM LatB and imaged while exposed at 20 °C.** Green represents LifeAct-GFP labeled F-actin and magenta the 3xmCherry-VAMP72A1 vesicles. Scale bar 10 μm.

**Movie S6: Representative myosin XIa TS 3xmCherry-VAMP72A1 cells treated with 0.25 μM LatB and imaged while exposed at 32 °C.** Green represents LifeAct-GFP labeled F-actin and magenta the 3xmCherry-VAMP72A1 vesicles. Scale bar 10 μm.

